# BIOLOGICAL SCIENCES: Genetics

**DOI:** 10.1101/404061

**Authors:** Jack S. Hsiao, Noelle D. Germain, Andrea Wilderman, Christopher Stoddard, Luke A. Wojenski, Geno J. Villafano, Leighton Core, Justin Cotney, Stormy J. Chamberlain

## Abstract

Angelman syndrome (AS) is a severe neurodevelopmental disorder caused by the loss of function from the maternal allele of *UBE3A*, a gene encoding an E3 ubiquitin ligase. *UBE3A* is only expressed from the maternally-inherited allele in mature human neurons due to tissue-specific genomic imprinting. Imprinted expression of *UBE3A* is restricted to neurons by expression of *UBE3A antisense transcript* (*UBE3A-ATS*) from the paternally-inherited allele, which silences the paternal allele of *UBE3A* in *cis*. However, the mechanism restricting *UBE3A-ATS* expression and *UBE3A* imprinting to neurons is not understood. We used CRISPR/Cas9-mediated genome editing to functionally define a bipartite boundary element critical for neuron-specific expression of *UBE3A-ATS* in humans. Removal of this element led to upregulation of *UBE3A-ATS* without repressing paternal *UBE3A*. However, increasing expression of *UBE3A-ATS* in the absence of the boundary element resulted in full repression of paternal *UBE3A*, demonstrating that *UBE3A* imprinting requires both the loss of function from the boundary element as well as upregulation of *UBE3A-ATS*. These results suggest that manipulation of the competition between *UBE3A-ATS* and *UBE3A* may provide a potential therapeutic approach for AS.

**SIGNIFICANCE STATEMENT:** Angelman syndrome is a neurodevelopmental disorder caused by loss of function from the maternal allele of UBE3A, an imprinted gene. The paternal allele of *UBE3A* is silenced by a long, non-coding antisense transcript in mature neurons. We have identified a boundary element that stops the transcription of the antisense transcript in human pluripotent stem cells, and thus restricts *UBE3A* imprinted expression to neurons. We further determined that *UBE3A* imprinting requires both the loss of the boundary function and sufficient expression of the antisense transcript to silence paternal *UBE3A*. These findings provide essential details about the mechanisms of *UBE3A* imprinting that may suggest additional therapeutic approaches for Angelman syndrome.

## Introduction

Angelman syndrome (AS) is a rare neurodevelopmental disorder characterized by developmental delay, seizures, lack of speech, ataxia, and severe intellectual disability (1, 2). It is most frequently caused by mutation (3, 4) or deletion (5) of the maternally inherited allele of *UBE3A*. *UBE3A* is an imprinted gene. The paternally inherited allele is silenced in brain (6, 7). The silencing of *UBE3A* is caused by expression of an opposing neuron-specific transcript antisense to *UBE3A* (*UBE3A-ATS)* (8). The regulation of *UBE3A-ATS* expression and the mechanism by which *UBE3A-ATS* represses *UBE3A* is of tremendous importance, since activation of paternal *UBE3A* is a promising therapeutic strategy for AS (9-11).

*UBE3A-ATS* is part of the >600 kb *SNURF/SNRPN* long non-coding RNA (heretofore referred to as *SNRPN*), which initiates from *SNRPN* promoters on the paternally inherited chromosome (12). The *SNRPN* lncRNA can be divided into two functional units based on tissue-specific transcription patterns in humans(13). The proximal portion of the transcript includes the protein-coding mRNAs, *SNURF* and *SNRPN*; two newly described long non-coding RNAs with snoRNA 5’ ends and polyadenylated 3’ ends, termed SPAs (14); snoLNC RNAs (15); the non-coding host gene for several C/D box small nucleolar RNAs (*SNORD109A, SNORD107, SNORD108*, and *SNORD116*); and the non-coding *IPW* transcript. The transcript containing these genes is ubiquitously transcribed in all tissues (16, 17). The distal portion of the transcript, which includes the non-coding host gene for additional small nucleolar RNAs (*SNORD115* and *SNORD109B*) and the non-coding *UBE3A-ATS*, is transcribed almost exclusively in the brain (13, 18-20). It is not known how the neuron-specific processing of *SNRPN* occurs, such that *UBE3A-ATS* expression, and thus *UBE3A* imprinting, is restricted to neurons.

We previously found that *UBE3A-ATS* was expressed and *UBE3A* was imprinted in non-neuronal cells derived from a patient with an atypical deletion of a portion of the paternal *SNRPN* allele (21). Based on these results, we hypothesized that imprinted expression of human *UBE3A* is restricted to neurons by a boundary element. Here we use CRISPR/Cas9 technology in human Angelman syndrome induced pluripotent stem cells (iPSCs) and their neuronal derivatives to functionally define this boundary element and determine its role in mediating *UBE3A* imprinting.

## Results

### A boundary element comprised of *IPW* and *PWAR1* restricts *UBE3A-ATS* expression to neurons

We previously reported that the distal portion of the *SNRPN* lncRNA is expressed and *UBE3A* is imprinted in induced pluripotent stem cells (iPSCs) derived from an individual with Prader-Willi syndrome (PWS) due to an atypical paternal deletion (21). This unique paternal deletion demonstrated that imprinting of *UBE3A* can occur in non-neuronal tissues and that a boundary may restrict expression of *UBE3A-ATS* and imprinting of *UBE3A* to neurons. The region separating the expressed proximal portion of the *SNRPN* lncRNA from the repressed distal portion includes a stretch of weak polyadenylation (poly(A)) sites at *IPW* (22) and two divergently-oriented CTCF binding sites at *PWAR1/PAR1 (*heretofore referred to as *PWAR1*; Fig. 1A; (23)). Poly(A) sites commonly mark the end of transcripts and signal transcriptional termination at the end of genes. CTCF is a structural protein with multiple potential functions, including insulating active and/or inactive chromatin domains and mediating long distance chromatin interactions. Publicly available RNA-seq data (www.encode.org; (24)) showed that most of the *SNRPN* lncRNA terminates at *IPW*, where the poly(A) sites are located, in most cell types. However, RNA polymerase II (RNAPII) was shown to accumulate further downstream within *PWAR1* in human embryonic stem cells (H1-ESC) (25)www.encode.org). These data led us to hypothesize that the two elements collectively efficiently terminate transcription of the *SNRPN* lncRNA in non-neuronal tissues (Fig 1A), thus restricting imprinted *UBE3A* expression to neurons.

**Figure 1.**
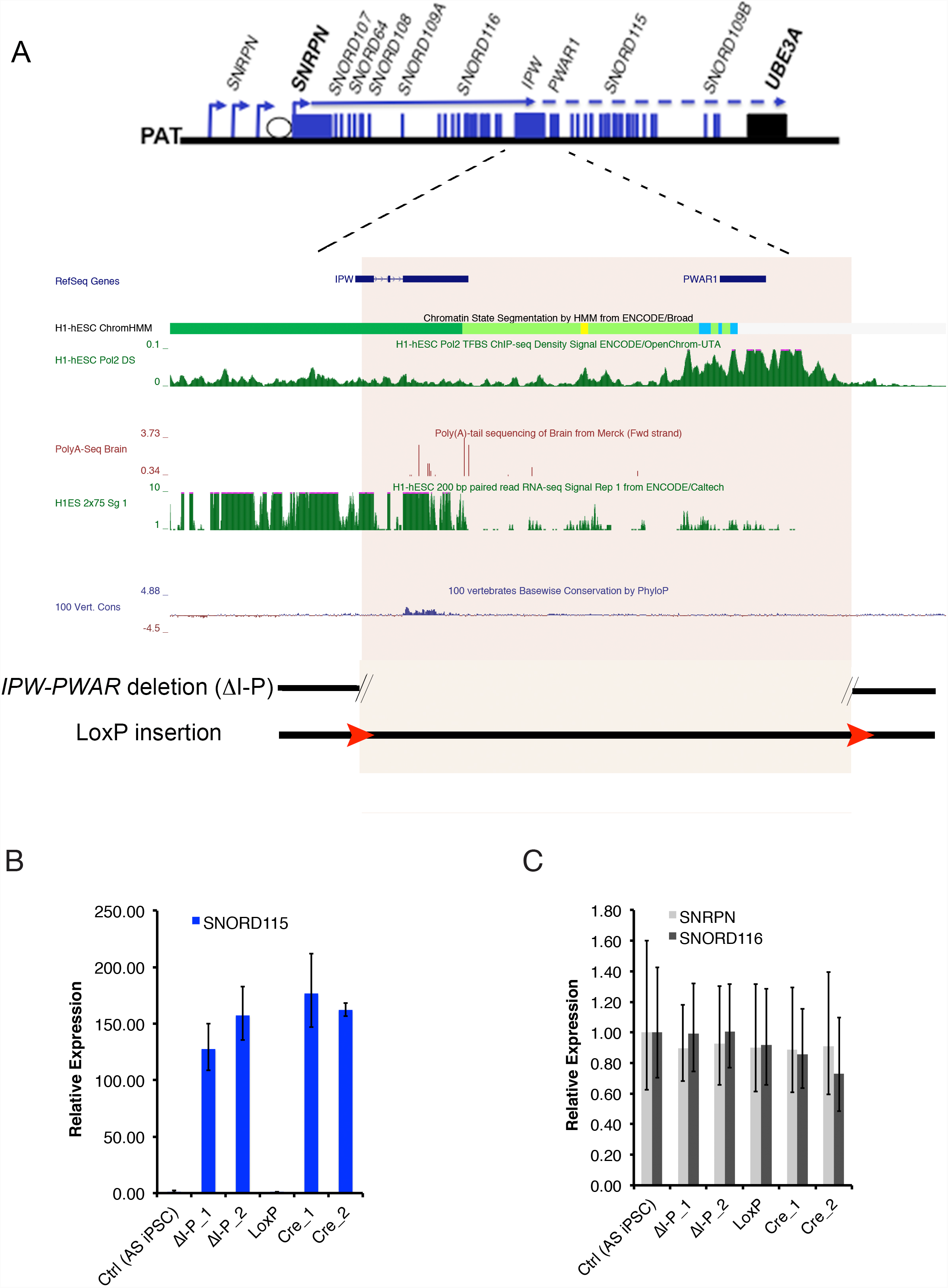
Deletion of a 24 kb region between *IPW* and *PWAR1* leads to ectopic expression of *SNORD115* in iPSCs. A) A diagram of the *SNRPN* transcriptional unit is shown (not to scale), followed by a more detailed view of the *IPW-PWAR1* region including UCSC Genome Browser data depicting genomic elements likely to contribute to the boundary function. Approximate deletion boundaries and loxP insertions are indicated at the bottom. B) RT-qPCR data quantifying *SNORD115* in iPSCs. C) RT-qPCR data quantifying *SNORD116* and *SNRPN* in iPSCs. δI-P indicates *IPW*-*PWAR1* deletion. _1 and _2 refer to independent clones generated using the same CRISPR constructs. LoxP indicates the floxed locus. Cre_1 and _2 are independent clones harboring the Cre-mediated deletion.

To test this hypothesis, we deleted a 24 kb region encompassing both *IPW* and *PWAR1* in AS iPSCs. These iPSCs harbor a ∼5.5 Mb deletion of the maternally-inherited allele of chromosome 15q11-q13, and thus enable us to easily focus on genes expressed from the paternal allele. A pair of CRISPRs designed to flank both *IPW* and *PWAR1* were electroporated into AS iPSCs along with two single stranded oligonucleotides (ssODNs) designed to insert LoxP sequences at the CRISPR cut sites following homology directed repair. After screening 96 clones using a PCR strategy modified from Kraft et al., we obtained 7 deletion clones and 1 clone with LoxP inserted at both cut sites. The LoxP sites were subsequently recombined using Cre-recombinase to create the 24 kb deletion. We also obtained 1 clone in which the sequence intervening the two CRISPR cut sites was inverted (INV). Two clones harboring CRISPR-mediated deletions of *IPW* and *PWAR1* (ΔI-P) and two clones from Cre-mediated recombination between LoxP sites (CreΔI-P) were chosen for further analysis. In iPSCs with both types of deletion, we detected the expression of *SNORD115* (Fig 1B), suggesting that the 24 kb region from *IPW* to *PWAR1* prevents expression of the distal portion of the *SNRPN* lncRNA in iPSCs. Deletion of this region did not affect the expression of the proximal coding and non-coding portions of *SNRPN* (Fig 1C).

### Both *IPW* and *PWAR1* contribute to boundary function

To decipher individual contributions of *IPW* and *PWAR1* to the boundary function, we deleted *PWAR1* (ΔP) and *IPW* (ΔI) separately in AS iPSCs (Fig 2A). In ΔP clones, we observed minimal expression of *SNORD115*. In ΔI clones, *SNORD115* expression was detected at approximately 50% of levels seen in ΔI-P clones. This suggested that the two components may work together to comprise full boundary function. Therefore, we deleted *IPW* and *PWAR1* sequentially, (ΔIΔP) leaving the sequence between the two elements intact. The expression levels of *SNORD115* in ΔIΔP clones was almost identical to those observed in ΔI-P clones (Fig 2A). This confirmed that *IPW* and *PWAR1* together are the pivotal elements providing boundary function between proximal and distal portions of the *SNRPN* lncRNA.

**Figure 2.**
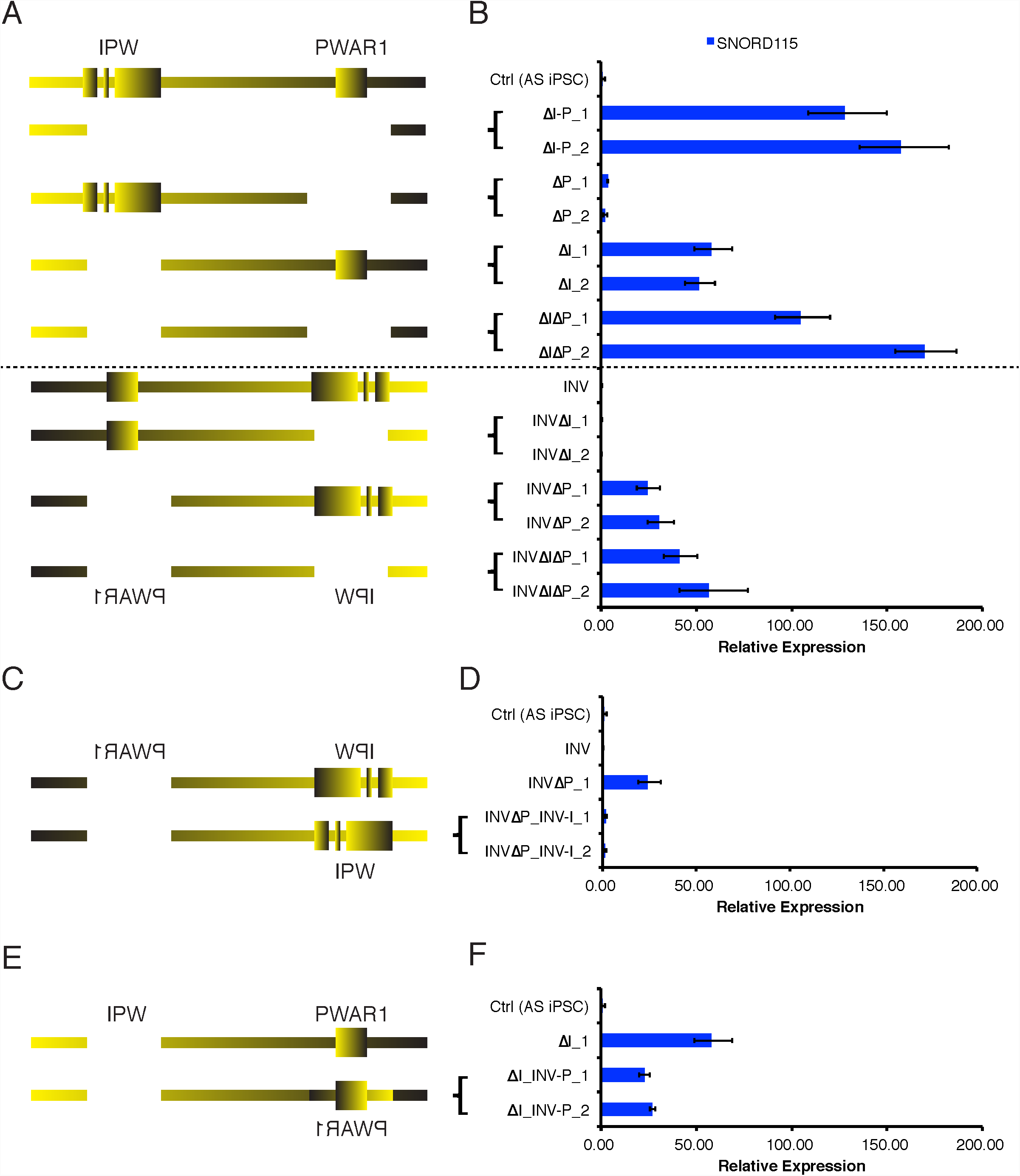
*IPW* and *PWAR1* both contribute to boundary function. Diagrams of CRISPR-mediated deletions/inversions generated in unmodified AS iPSCs (A), INV AS iPSCs (C), and ΔI AS iPSCs (D) are shown. RT-qPCR for S*NORD115* in iPSCs with the corresponding deletion/inversion is shown in B, D, and F. _1 and _2 denote two independent clones generated using the same CRISPR constructs.

Poly(A)-dependent transcriptional termination requires proper orientation of the poly(A) sequence and downstream sequences required to bind cleavage stimulation factor and enhance poly(A)-dependent cleavage (26). Recent studies also suggest that the orientation of CTCF can influence its ability to form chromatin loops, although presumably, not all functions of CTCF require a specific orientation (27, 28). Paradoxically, when we inverted the 24kb boundary in AS iPSCs (INV; Fig 2B), we did not detect *SNORD115* expression, suggesting that the boundary was still functional in the inverted orientation. To further understand this paradox, we deleted *IPW* and *PWAR1* separately in the INV iPSCs. We did not detect *SNORD115* when *IPW* was deleted in INV iPSCs (INVΔI; Fig 2B). However, when *PWAR1* was deleted in the INV iPSCs, *SNORD115* was detected (INVΔP; Fig 2B). Notably, *SNORD115* expression in INVΔP lines is about 40% of that in ΔI-P lines (Fig 2A). Sequential deletion of *IPW* and PWAR1 in the INV iPSCs (INVΔPΔI) resulted in a slight increase in *SNORD115* expression, but did not fully restore expression to the levels seen in ΔI-P or ΔIΔP iPSCs.

We took advantage of the fact that *SNORD115* is expressed in ΔI and INVΔP iPSCs to individually test the directionality of *IPW* and *PWAR1*. We first restored *IPW* to its natural orientation in INVΔP iPSCs and found that *SNORD115* expression was barely detectable (INVΔP_INV-I; Fig 2C), demonstrating that *IPW* can stop transcription in its natural orientation. Next, we inverted *PWAR1* in ΔI iPSCs, and found that *SNORD115* expression was reduced by 50% compared to the ΔI parent line, suggesting that the inverted *PWAR1* gained a new function ((ΔI_INV-P); Fig 2D). Together, these results suggested that both elements within the boundary require proper orientation to function appropriately.

### Long-distance interactions involving *IPW* and *PWAR1*

*IPW* and *PWAR1* constitute a strong chromatin boundary that may coincide with a putative topologically associated domain (TAD), based on published Hi-C data (29). To determine whether boundary function involves specific 3D interactions, we first asked whether CTCF is bound to the *PWAR1* region. CTCF is a structural protein that mediates chromatin loops and can separate chromatin boundaries. *PWAR1* hosts a cluster of two divergent CTCF binding sites. We performed ChIP-seq and ChIP-qPCR using antibodies against CTCF in iPSCs and iPSC-derived neurons with large deletions of maternal and paternal chromosome 15q11-q13. CTCF was bound at several sites across the imprinted domain on the paternally-inherited allele in AS iPSCs, including the *PWAR1* exon (Fig S1A). However, the entire imprinted domain was largely devoid of CTCF binding in PWS iPSCs, which carry only a maternal allele of chromosome 15q11-q13 (Fig S1A). We identified allele-specific binding of CTCF at nine sites across the imprinted domain in iPSCs (Table Sx). CTCF binding outside of the imprinted domain was nearly identical in AS and PWS iPSCs (Fig S1A). Upon differentiation of AS iPSCs into neurons, CTCF binding at *PWAR1* as well as several other sites was reduced (Fig S1C,D). We observed retained CTCF binding in neurons at two different sites on the paternal allele, however (Fig S1B). CTCF binding at sites upstream of *SNRPN* and *UBE3A* promoters remained intact during the 10-week time course of neural differentiation.

Next, we utilized circularized chromosome conformation capture followed by sequencing (4C-seq) to determine whether *IPW* and *PWAR1* relied on specific long distance interactions to confer boundary function. 4C enables the identification of all loci that interact with a specific viewpoint of choice. We performed 4C-seq using viewpoints located at *IPW* and *PWAR1* in AS iPSCs and 10-week neurons (Fig 3). In iPSCs, the *IPW* viewpoint only showed significant interactions with *PWAR1* and points upstream of it. In neurons, *IPW* interactions were mapped to points upstream and downstream, including the *UBE3A* promoter. Thus, *IPW* does not interact across the boundary in iPSCs, but does in neurons, where the boundary is dissolved. The CTCF binding sites at *PWAR1* showed significant interactions with points upstream and downstream of the boundary in iPSCs. Points upstream that interact with the CTCF sites at *PWAR1* include the upstream exons of *SNRPN*, which are annotated as strong enhancer or promoter states. Points downstream interacting with *PWAR1* in iPSCs include a CTCF site at the distal end of *SNORD115*. In neurons, *PWAR1* has few interactions and they are local. These data demonstrate that the 24kb boundary restricts 3D interactions with *IPW* in iPSCs. Although, 3D interactions with the CTCF sites at *PWAR1* differ between iPSCs and neurons, they do not seem to be restricted by boundary function. In fact, 3D interactions with *PWAR1* in iPSCs are more consistent with an interaction between the alternative upstream promoters of *SNRPN* and the 3’ end of transcripts originating there.

**Figure 3.**
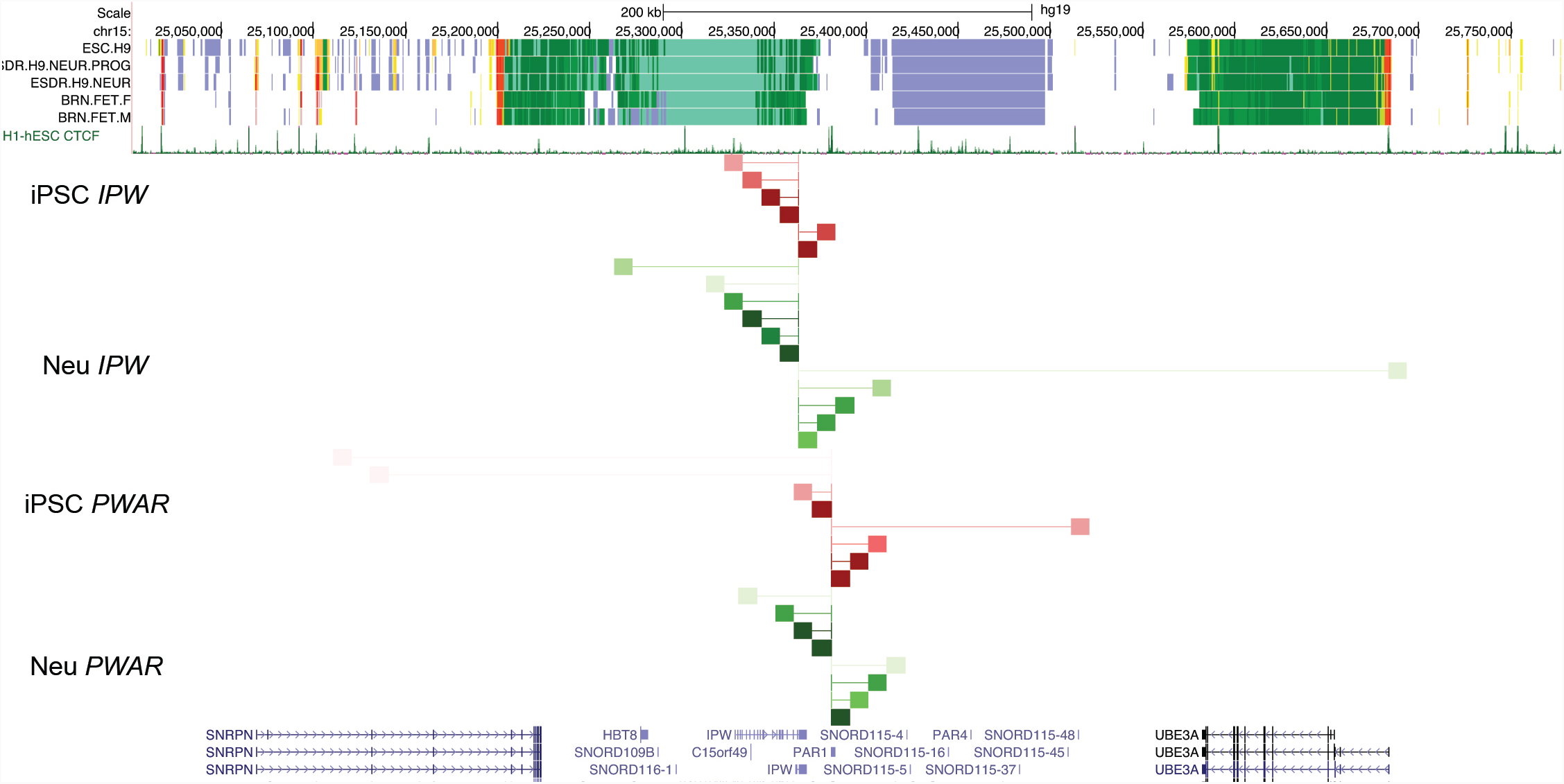
3D interactions with *IPW* and *PWAR1*. Analysis of 4C-seq data are shown, along with chromatin state annotations from H9 hESCs, H9-derived neural progenitors, H9-derived neurons, and male/female fetal brain tissues from the Roadmap Epigenomics Project. CTCF binding sites and UCSC genes are shown for reference. Red lines and blocks refer to interactions in AS iPSCs, green lines and blocks refer to interactions in AS iPSC-derived neurons. The thin vertical lines at *IPW* and *PWAR1* refer to the anchor point for 4C-seq. All interactions are significant (p<0.001) with darker colors indicating decreased p-value (higher significance).

### *UBE3A* imprinting requires sufficient levels of *UBE3A-ATS* expression

We previously reported imprinted *UBE3A* expression in an iPSC line that aberrantly expresses *UBE3A-ATS* due to an atypical PWS deletion. Based on these data, we predicted that *UBE3A* would be imprinted in iPSCs expressing *SNORD115* and *UBE3A-ATS.* To our surprise, *UBE3A* imprinting was not observed in ΔI and ΔI-P clones where *UBE3A-ATS* is transcribed (Fig 4B). Therefore, we tried to recapitulate our previous observation with the atypical PWS deletion in an AS iPSC line(21). We used CRISPR/Cas9 to remove a 303 kb region between *SNRPN* intron 1 and the last copy of *SNORD115* (*SNORD115*-47) in AS iPSCs (ΔS-115; Fig 4A). This deletion juxtaposes the *SNRPN* promoter(s) immediately upstream of *UBE3A-ATS*. Indeed, paternal *UBE3A* is completely repressed in iPSCs with this deletion (Fig 4C) suggesting that increasing *UBE3A-ATS* transcription is necessary to imprint *UBE3A*.

**Figure 4.**
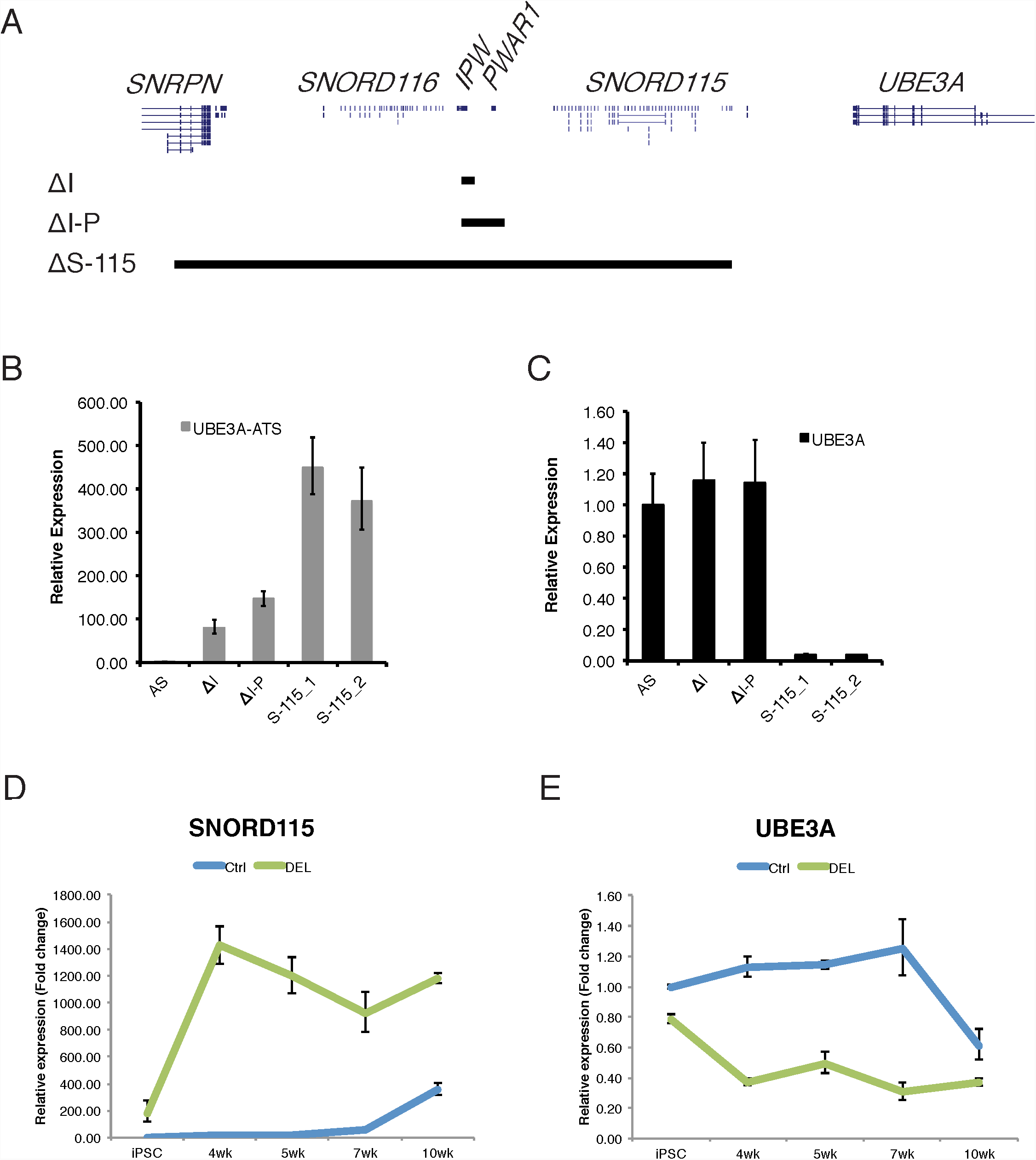
Sufficient expression of *UBE3A-ATS* is required to imprint *UBE3A*. Diagram depicting relative sizes of ΔI, ΔI-P, and ΔS-115 deletions is shown in A. RT-qPCR for *UBE3A-ATS* and *UBE3A* are shown in B and C, respectively. RT-qPCR for *SNORD115* and *UBE3A* is shown across a time course of neural development in AS iPSCs (Ctrl) and ΔI-P AS iPSCs (DEL) in D and E, respectively.

Since transcription of the *SNRPN* lncRNA is normally increased during neurogenesis, we sought to determine whether an early increase in expression of *UBE3A-ATS* during neurogenesis would lead to premature imprinted *UBE3A* expression in neural derivatives of ΔI-P iPSCs, which lack the boundary. We differentiated AS and ΔI-P iPSCs into forebrain cortical neurons as previously described (30) and collected RNA samples during the time course of differentiation. We found that *SNORD115* expression increases and *UBE3A* becomes silenced between weeks 7 and 10 of differentiation in AS iPSCs, consistent with our previously published observations (Fig 4D,E; (13, 18)). The ΔI-P iPSCs showed a slight reduction of *UBE3A* expression compared to AS iPSCs. By 4 weeks of neural differentiation, *SNORD115* expression in ΔI-P neural progenitors is increased to maximum levels and *UBE3A* attains it lowest expression levels (Fig 4D,E). These data demonstrate that sufficient levels of *UBE3A-ATS* transcription are necessary to silence *UBE3A,* and that the 24 kb boundary element also regulates the timing of *UBE3A* imprinting during neurogenesis.

*UBE3A-ATS* is expressed in ΔI and ΔI-P iPSCs, but *UBE3A* is not imprinted. On the other hand, *UBE3A-ATS* is expressed and *UBE3A* is imprinted in ΔS-115 iPSCs, enabling us to study AS iPSCs that imprint and do not imprint *UBE3A*. We sought to visualize and compare the interactions between *UBE3A-ATS* and *UBE3A* under these conditions. We performed precision nuclear run-on sequencing (PRO-seq) on these samples. PRO-seq determines the active sites of transcriptionally engaged RNAPII by mapping nascent transcription (31, 32). PRO-seq data from iPSC lines revealed plus-strand RNAPII density across *UBE3A-ATS* in ΔI, ΔI-P, and ΔS-115 iPSCs (Fig 5). Minus strand RNAPII density was seen across the entire *UBE3A* gene in all iPSCs, but the ΔS-115 iPSCs had robust PRO-seq density only in the first half of the gene (Fig 5). These data suggest *UBE3A* imprinting coincides with reduction of full-length transcript, since polymerases do not appear to efficiently make it to the 3’ end of the gene.

**Figure 5.**
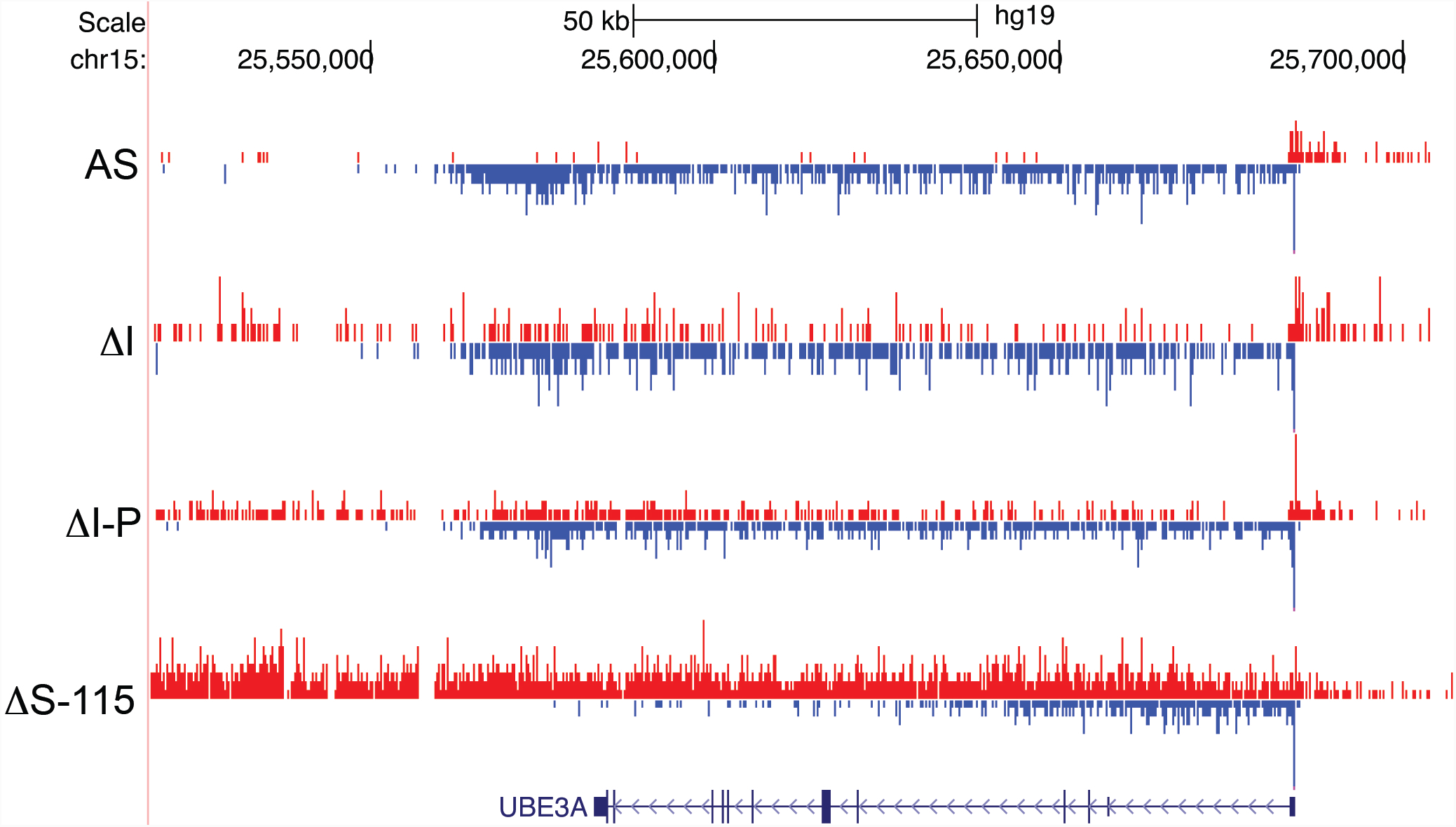
Imprinting of *UBE3A* coincides with reduced RNAPII density across 3’ half of *UBE3A* gene body. PRO-seq was used to map RNAPII density in AS, ΔI, ΔI-P, and ΔS-115 iPSCs. Plus strand RNAPII density is shown in red. Minus strand RNAPII density is shown in blue.

## Discussion

Imprinted expression of *UBE3A* is restricted to neurons by the tissue-specific expression of *UBE3A-ATS* (11, 33, 34). *UBE3A-ATS* is at the 3’ end of the *SNRPN* lncRNA, which also includes host genes for *SNORD116* and *SNORD115*, as well as other non-coding RNAs(12). In humans, the proximal half of the *SNRPN* lncRNA is expressed broadly in different tissue types, while the distal half, including *UBE3A-ATS* is restricted to neurons (13, 18, 19). We used CRISPR/Cas9 to functionally define the boundary element that restricts *UBE3A-ATS* expression to neurons (Fig 2). We found the boundary to be comprised of two parts: one part includes poly(A) and conserved sequences in the last exon of *IPW*, while the other includes a cluster of CTCF sites in and around the exon annotated as *PWAR1*. Although both elements contribute to boundary function, *IPW* plays a larger role, and is required to completely stop transcription in non-neuronal cells. *IPW* requires its natural orientation to stop transcription, suggesting that the poly(A) sites are important for boundary function. CTCF binds to the *PWAR1* exon in iPSCs, but not in neurons, suggesting that CTCF binding may contribute to boundary function as well. PRO-seq experiments demonstrate reduced RNAPII density downstream of *PWAR1* in ΔI iPSCs, suggesting that these CTCF sites may pause RNAPII and facilitate RNAPII disengagement (Suppl. Fig 2). CTCF has been previously shown to pause elongating RNAPII to influence alternative splicing (35). Interestingly, RNAPII is paused and/or disengaged near the first exons encoding *SNORD115* in ΔI-P iPSCs by an as-yet-unknown mechanism. This suggests multiple redundancies may prevent *UBE3A* imprinting in this cell type. Based on these findings, we propose a simple model by which this bipartite boundary element stops transcription in most cell types. We propose that the poly(A) sites within *IPW* stop transcription via poly(A)-dependent cleavage, while CTCF binding at *PWAR1* slows RNAPII enough to allow the XRN2 5’-3’ exonuclease to lead to termination in what is known as the ‘torpedo model’ of transcription termination(36, 37).

It is not clear how the boundary function is lost during neurogenesis. 4C-seq experiments demonstrate that 3D interactions with *IPW*, are restricted to sites upstream in iPSCs, but are bi-directional in neurons, consistent with a loss of boundary function during neurogenesis. CTCF binding within *PWAR1* is present on the paternal allele in iPSCs, but not in neurons. This loss of CTCF binding may contribute to loss of boundary function in neurons. Consistent with this hypothesis, sites interacting with *PWAR1* in neurons are limited to nearby loci, are largely not bound by CTCF, and overlap with several sites interacting with *IPW*.

We further speculate that loss of CTCF binding may contribute to reduced termination at *IPW* in neurons. CTCF is gradually lost from *PWAR1* during the 10-week course of neural differentiation (Fig S1C), correlating with full expression of *UBE3A-ATS* and imprinting of *UBE3A* (Fig 4D,E). iPSCs lacking the bipartite boundary imprint *UBE3A* precociously during neuronal differentiation, supporting the hypothesis that the boundary element also controls the developmental timing of *UBE3A* imprinted expression. An understanding of how *IPW* and *PWAR1* independently contribute to the developmental timing of *UBE3A* imprinting may help determine how they facilitate boundary removal during neurogenesis. Paradoxically, deletion of *PWAR1*--including both CTCF sites—does not substantially decrease transcriptional termination in iPSCs (Fig 1A). Perhaps this is due to the presence of additional elements capable of pausing RNAPII. Indeed PRO-seq data reveal RNAPII pausing near the first exon of the *SNORD115* cluster (Fig S2).

Finally, the surprising observation that *UBE3A-ATS* is expressed, but *UBE3A* is not imprinted in iPSCs with deletions of *IPW* or *IPW* plus *PWAR1* (Fig 4C) indicate that imprinted expression of *UBE3A* also requires sufficient expression of *UBE3A-ATS*, in addition to loss of boundary function. Indeed, a CRISPR-mediated deletion that increases *UBE3A-ATS* expression led to full repression of paternal *UBE3A*. PRO-seq experiments further demonstrated that *UBE3A* imprinting in these iPSCs coincided with reduced active RNAPII across the 3’ half of *UBE3A* (Fig 5). These data further support the notion that *UBE3A-ATS* represses paternal *UBE3A* via transcriptional interference. If *UBE3A* imprinting occurs due to transcriptional interference, manipulation of *UBE3A-ATS* <u>or</u> *UBE3A* transcription may provide alternative therapeutic approaches for Angelman syndrome.

## Materials and Methods

### Cell Culture

AS iPSC (AS del 1-0) and PWS iPSC (PWS del 1-7) lines were generated and maintained by mechanical passaging on mouse embryonic fibroblasts (MEFs) as previously described(13, 18).

### CRISPR genome editing

CRISPR gRNA sequences were designed using CRISPR Genome Engineering Resource (http://crispr.mit.edu)(39), and cloned into the px459 V2 vector(40, 41). The sequence of CRISPRs and ssODNs used in this paper are listed in Table 2. To introduce the CRISPRs into AS iPSCs, 10 μg of each of two CRISPRs flanking the region to be deleted were electroporated into 6-10 million AS iPSCs that were treated with 10 μM ROCKi (Calbiochem; Y-27632)(42, 43) 24 hours prior to electroporation. Cells were then seeded on DR4 MEFs and treated with ROCKi for 48 hours. Positive selection for the presence of the CRISPRs was then carried out for 48 hours with 1 μg/ml puromycin beginning 24 hours after seeding on MEFs.

### ChIP

ChIP qPCR was performed using Millipore EZ-Magna ChIP G – Chromatin Immunoprecipitation Kit (17-409) following manufacturer’s instructions using 6-10 million cells. ChIPAb+ CTCF (17-10044) antibody was used to immunoprecipitate sonicated chromatin overnight. SYBR green primers (Table 1) were used for ChIP-qPCR. For ChIP-seq, library preparation and sequencing was performed by the Genomics Core in the Yale Stem Cell Center (https://medicine.yale.edu/stemcell/coreservices/corelabs/genomics.aspx). FASTQ files were mapped and analyzed using Homer with the parameters described previously(44, 45).

### 4C-seq

4C-seq was carried out as described(46) using nuclei harvested from approximately 2 million formaldehyde-fixed and glycine-quenched iPSCs and iPSC-derived neurons. NlaIII enzyme was used for the first digestion, and DpnII was used for the second digestion. Data were analyzed using the r3Cseq package(47).

### PRO-seq

PRO-seq was carried out as described(31, 32, 48) using 1 x 10^6^ permeabilized cells per iPSC line.

Detailed Materials and Methods are found in Supplementary information.

## Acknowledgements

The authors would like to thank Ms. Carissa Sirois and Dr. Marc Lalande for helpful discussions. The work was supported by the following funding sources: NIH R01HD068730, Angelman Syndrome Foundation, and Connecticut DPH Stem Cell Research Program (12SCBUCHC) to SJC and NIH R35GM119465 to JC.

---

**Suppl. Fig. 1. CTCF is enriched on the paternal allele of *PWAR1.*** ChIP-seq for CTCF was performed in AS and PWS iPSCs harboring deletion of the 15q11-q13 region on the maternal and paternal alleles, respectively. CTCF binding on the paternal allele (blue) and maternal allele (red) are shown in A. The imprinted region is designated. Asterisks mark the CTCF sites used as anchors for 4C-seq. Black bars indicate significant differentially bound CTCF sites based on ChIP-Seq data. ChIP-qPCR was used to quantify CTCF enrichment on the paternal and maternal alleles in AS and PWS iPSCs, respectively, in B. ChIP-qPCR was used to quantify CTCF enrichment at *PWAR1* and *SNORD116* sites (C) and at sites upstream of SNRPN and upstream of *UBE3A* (D) during a time course of neural differentiation of AS iPSCs.

**Suppl. Fig. 2. RNAPII is stalled at *PWAR1* and at the first exon of *SNORD115* in AS iPSCs.** PRO-seq was used to map RNAPII density in AS, ΔI, ΔI-P, and ΔS-115 iPSCs. Plus strand RNAPII density is shown in red. Minus strand RNAPII density is shown in blue. Absent signals in ΔI, ΔI-P, and ΔS-115 iPSC lanes reflect engineered deletions.

